# An annotated chromosome-scale reference genome for Eastern Black-eared Wheatear (*Oenanthe melanoleuca*)

**DOI:** 10.1101/2022.12.22.521689

**Authors:** Valentina Peona, Octavio Manuel Palacios-Gimenez, Dave Lutgen, Remi André Olsen, Niloofar Alaei Kakhki, Pavlos Andriopoulos, Vasileios Bontzorlos, Manuel Schweizer, Alexander Suh, Reto Burri

**Author notes:** These authors contributed equally to the work (Valentina Peona, Octavio Manuel Palacios-Gimenez, Dave Lutgen, Reto Burri). **Correspondence** Reto, Swiss Ornithological Institute, Seerose 1, CH-6204 Sempach.

## Abstract

Pervasive convergent evolution and in part high incidences of hybridization distinguish wheatears (songbirds of the genus *Oenanthe*) as a versatile system to address questions at the forefront of research on the molecular bases of phenotypic and species diversification. To prepare the genomic resources for this venture, we here generated and annotated a chromosome-scale assembly of the Eastern Black-eared Wheatear (*O. melanoleuca*). This species is part of the *O. hispanica-*complex that is characterized by convergent evolution of plumage coloration and high rates of hybridization. The long-read-based male nuclear genome assembly comprises 1.04 Gb in 32 autosomes, the Z chromosome, and the mitogenome. The assembly is highly contiguous (contig N50: 12.6 Mb; scaffold N50: 70 Mb), with 96% of the genome assembled at chromosome level and 96% BUSCO completeness. The nuclear genome was annotated with 18’160 protein-coding genes and 30’730 mRNAs, and about 10% of the genome consists of repetitive DNA. The annotated chromosome-scale reference genome of Eastern Black-eared Wheatear provides a crucial resource for research into the genomics of adaptation and speciation in an intriguing group of passerines.

## Introduction

Wheatears of the genus *Oenanthe* and their relatives – together referred to as “open-habitat chats” – are a group of songbirds that display several remarkable characteristics distinguishing them as a versatile system to address key questions on the evolution of phenotypes and formation of species. Many phenotypes, including multiple conspicuous colour ornaments, seasonal migration, and sexual dimorphism appear independently in multiple branches within open-habitat chats, suggesting a high incidence of convergent evolution (Alaei Kakhki et al. in press; Aliabadian et al. 2012; Schweizer et al. 2019). Furthermore, hybridization is observed in several species complexes and occurs at notably high rates in the *O. hispanica-*complex that consists of four currently recognized taxa (Schweizer et al. 2019): Western Black-eared Wheatear (*O. hispanica*), Pied Wheatear (*O. pleschanka*), Cyprus Wheatear (*O. cypriaca*), and Eastern Black-eared Wheatear (*O. melanoleuca*; **Fig. 1**). Pied and Eastern Black-eared Wheatear hybridize pervasively at the western shores of the Black Sea, in the Caucasus, and in the Alborz mountains of northern Iran (Haffer 1977; Panov 2005). The resulting introgression reaches beyond the hybrid zones (Schweizer et al. 2019), and hybrid zones themselves sport admixed phenotypes that display combinations of plumage colour phenotypes divergent between species (mantle and neck-side coloration) (Haffer 1977; Panov 2005). Finally, a phenotype divergently expressed between many wheatear species, black-or-white throat coloration, segregates as polymorphisms in three species of the *O. hispanica-*complex. Once a high-quality reference genome is available, this polymorphism and the recombination of mantle and neck-side coloration in hybrids provide an excellent opportunity to map these phenotypes to the genome (Buerkle and Lexer 2008) and study their convergent evolution across open-habitat chats. Furthermore, hybridization in several geographic regions enables insights into common or idiosyncratic patterns of evolution under hybridization (Gompert et al. 2017).

**Figure 1.**
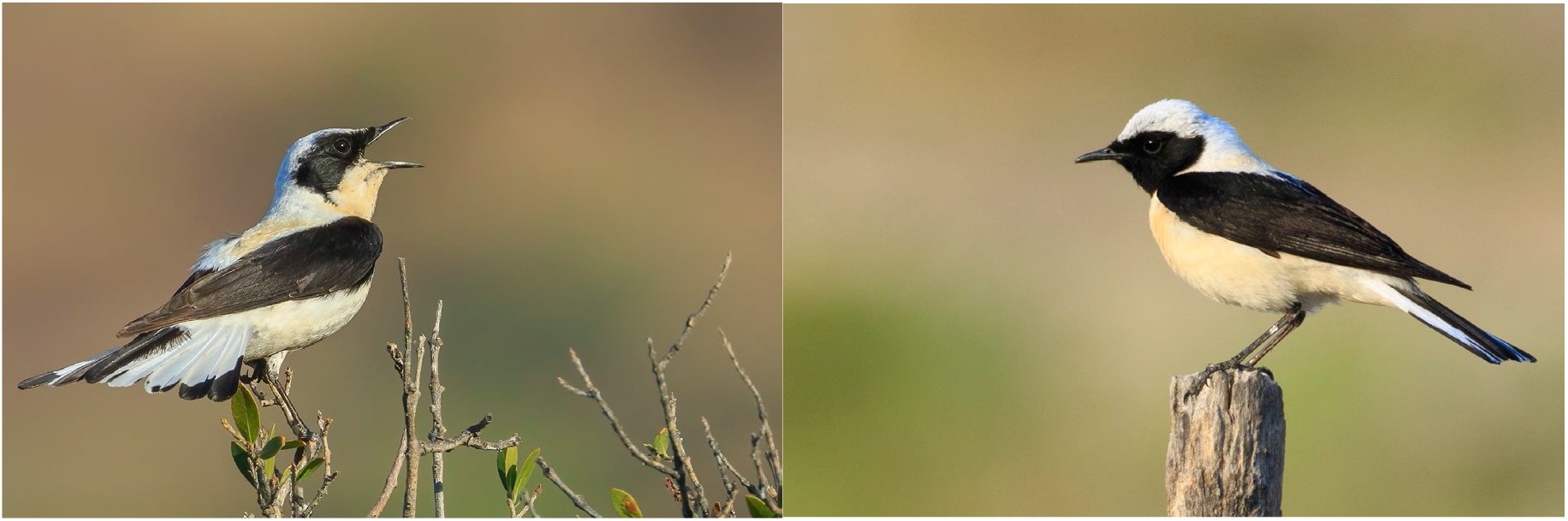
Eastern Black-eared Wheatear (*Oenanthe melanoleuca*). The species sports a white-throated (left; Agii Pantes, Greece, June 2022) and a black-throated phenotype (right; Lesvos, Greece, May 2017) in males. © Reto Burri

Here, we describe the *de novo* assembly and annotation of a chromosome-scale reference genome for the Eastern Black-eared Wheatear (*O. melanoleuca*). The assembly includes models for 32 autosomes, the Z chromosome and the mitogenome that together cover 90% of the k-mer-based genome size estimate (94% with unplaced scaffolds included); it is highly contiguous with a scaffold N50 of 70 Mb and BUSCO completeness score of 96%. This reference genome enables genomic research into the evolutionary history of phenotypic and species diversification in wheatears and their close relatives.

## Material and Methods

### Sampling, tissue preservation, and nucleic acid extraction

To obtain optimal starting material for a reference individual, we freshly sampled a male Eastern Black-eared Wheatear (*Oenanthe melanoleuca*) in Galaxidi, Greece (sampling permit no. 181968/989, issued by the Ministry of Environment and Energy, General Secretariat of Environment, General Directorate of Forests and Forest Environment, Directorate of Forest Management, Department of Wildlife and Game Management; export permit no. 55980/1575, Regional CITES management authority Attika). For this purpose, we sampled about 100 μl of blood from the brachial vein, and, after euthanizing the bird, we extracted all tissues possible. Tissues were immediately snap-frozen in liquid nitrogen. Throughout transportation and storage preceding DNA extraction, the samples were kept at a temperature below -80° C.

To obtain ultra-high molecular weight (UHMW) DNA from the reference individual, NGI Uppsala (Sweden) extracted DNA from the blood sample using the Bionano Prep™ Blood and Cell Culture DNA Isolation Kit (Bionano, San Diego, USA). Electrophoresis on a Femto Pulse instrument showed a mean DNA fragment length of about 200 kb, with fragments reaching up to 800 kb.

To prepare muscle tissue for Hi-C sequencing library preparation, we pulverized breast muscle tissue from the reference individual in a mortar. To avoid unfreezing of the tissue powder, the procedure was carried out in a climate chamber at 4°C under regular addition of liquid nitrogen.

To prepare RNA for full-length transcript sequencing, we extracted total RNA from eight snap-frozen tissues kept at -80°C (brain, breast muscle, heart, kidney, liver, lung, spleen, and testis) using the RNeasy Mini Kit (Qiagen; Hombrechtikon, Switzerland) according to the manufacturer’s instructions. RNA quality was assessed with a Fragment Analyzer (Agilent). RNA from spleen showed considerable degradation and was excluded from further analyses.

### *De novo* genome sequencing, and reference genome assembly and annotation

#### Assembly strategy and data acquisition

To obtain a chromosome-scale reference genome, our strategy largely followed the multiplatform approach recommended by Peona et al. (2021). In brief, it consisted of (i) a phased primary assembly based on long reads (ii) polishing and scaffolding of the primary assembly with linked-read sequencing data, and (iii) scaffolding of the secondary assembly with proximity ligation (Hi-C) information.

To this end, we obtained a total of 215 Gb (unique coverage 151 Gb) Pacific Biosciences (PacBio) long-read sequence data, 54 Gb linked-read sequence data, and 83 Gb Hi-C data. NGI Uppsala (Sweden) prepared a PacBio library from UHMW DNA and sequenced this library on 18 SMRT Cells 1M on a PacBio Sequel instrument. A linked-read sequencing library was prepared using the 10X Genomics Chromium Genomic Kit (from the same DNA extraction as used for PacBio sequencing; 10X Genomics, Inc., Pleasanton, CA, USA; Cat No. 120215), and a Hi-C library was prepared with the Dovetail Omni-C kit (Scotts Valley, CA, USA; Cat No. 21005). The linked-read and Hi-C libraries were prepared and sequenced on a NovaSeq 6000 instrument (S4 lane, 150 bp paired-end reads) at the facilities of NGI Stockholm (Sweden).

#### Genome size estimation

We estimated genome size by counting k-mer frequency of the quality-checked 10X Genomics linked reads. To this end, we first trimmed 22 bp from all 10X Genomics linked reads using fastp (Chen et al. 2018) to remove indices from R1 reads and keep symmetric read lengths for the R2 reads. We then counted k-mers of size 21 using jellyfish 2.2.10 (Marçais and Kingsford 2011) and used GenomeScope (Vurture et al. 2017) to estimate genome size from k-mer count histograms.

#### *De novo* genome assembly

We assembled the PacBio long reads into the phased primary assembly using the Falcon Unzip assembler (Chin et al. 2016), followed by polishing with Arrow. Before assembly polishing, we masked repeat regions of the phased primary assembly with RepeatMasker (Smit et al. 1996-2010) using a custom repeat library (Boman et al. 2019; Peona et al. 2021; Suh et al. 2018; Weissensteiner et al. 2020) to make accurate assembly corrections without overcorrecting large repeats. We then polished the masked assembly with two rounds in Pilon v1.22 (Walker et al. 2014) with the parameter “--fix indels” using the reference individual’s linked-read data. To purge duplicate scaffolds from the assembly, we ran purge_dups (Guan et al. 2020) on the polished assembly. Prior to scaffolding with linked-read data, we split potential mis-assemblies with reference-individual linked-read data using Tigmint (Jackman et al. 2018). With the aim to scaffold the polished remaining contigs, we applied ARCS+LINKS using the reference individual’s linked-read data using default parameters (Warren et al. 2015; Yeo et al. 2018).

To further scaffold the assembly, we applied the 3D-DNA pipeline (Dudchenko et al. 2017) to join the sequences into chromosomes. We first used Juicer v.1.6 (Durand et al. 2016) to map Hi-C data against the contigs and to filter reads, and then ran the asm-pipeline v.180922 to generate a draft scaffolding.

Finally, we corrected mis-assemblies based on the visual inspection of the proximity map using Juicebox (Robinson et al. 2018). The final chromosome-level assembly was polished with two additional rounds in Pilon as described above.

To assess homology of the assembled scaffolds with bird chromosomes, we aligned the final genome assembly to the genomes of collared flycatcher (*Ficedula albicollis*) (FicAlb1.5) (Kawakami et al. 2014), zebra finch (taeGut3.2.4) (Warren et al. 2010), and chicken (GRCg6a) (Bellott et al. 2017) using D-Genies (Cabanettes and Klopp 2018). Chromosomes were named according to homology with the latter three genomes. In cases, such as chicken chromosomes 1 and 4 that are split to multiple chromosomes in songbirds, the nomenclature in the wheatear genome was adapted to the species whose homologous chromosome matched closest.

#### Mitogenome assembly

To assemble the mitochondrial genome, we used the MitoFinder 1.4 (Allio et al. 2020) and mitoVGP 2.2 (Formenti et al. 2021) pipelines with the published *Oenanthe isabellina* mitochondrial genome (Genbank Accession Number: NC_040290.1) as reference. We ran MitoFinder with the reference individual’s short-read data (linked-read data but without making use of the linked-read haplotype information), and with mitoVGP we made joint use of the linked-read and long-read data. From MitoFinder we extracted the longest contig containing all 13 protein coding genes, two rRNA genes and 22 tRNAs annotated by MitoFinder as mitogenome assembly. We annotated both assemblies using the MITOS WebServer (http://mitos2.bioinf.uni-leipzig.de/index.py).

We then aligned both resulting assemblies with the mitogenomes of Isabelline Wheatear (*O. isabelline*, NC_040290.1) and Northern Wheatear (*O. Oenanthe*, MN356231.1) using MUSCLE (Edgar 2004) in MEGA X (Stecher et al. 2020) and generated a circular mitogenome map using CGView (Stothard and Wishart 2005).

#### Assembly quality evaluation

To evaluate assembly quality at each assembly step, we estimated basic assembly statistics using QUAST (Gurevich et al. 2013) and evaluated the completeness of expected gene content in the assembly based on benchmarking universal single-copy orthologs (BUSCO) (Simão et al. 2015) with the avian dataset aves_odb10 (8’338 BUSCO) in BUSCO 5.0.0.

#### Repeat annotation

The final version of the genome assembly was used to *de novo* characterize both interspersed and tandem repeats. For interspersed repeats, we used RepeatModeler2 (Flynn et al. 2020) with the option “-LTR_struct” to obtain an improved characterisation of LTR retrotransposons which are commonly found in avian genomes (Boman et al. 2019; Kapusta and Suh 2017; Peona et al. 2021). The resulting library of raw consensus sequences was filtered from consensus sequences of tandem repeats (for which we ran a specific analysis; see below) and from protein-coding genes using the Snakemake pipelinerepeatlib_filtering_workflow v0.1.0 (https://github.com/NBISweden/repeatlib_filtering_workflow).

For tandem repeats, we used RepeatExplorer2 (Novák et al. 2020) to search for satellite DNA (satDNA) sequences using the reference individual’s 10X Genomics linked reads. Prior to RepeatExplorer2 graph-based clustering analysis, sequencing reads were pre-processed and checked by quality with FastQC (Babraham Bioinformatics: Cambridge 2012) using the public online platform at https://repeatexplorer.elixir-cerit-sc.cz. We processed the reads with the “quality trimming tool”, “FASTQ interlacer on the paired end reads”, “FASTQ to FASTQ converter”, followed by “RepeatExplorer2 clustering” with default parameters. Each reference sequence assembled by RepeatExplorer2 consisted of a monomer of the satDNA consensus sequence. The relative genomic abundance and nucleotide divergence (Kimura-2-parameter distance) of each detected satDNA were estimated by sampling four million read pairs and aligning them to the satDNA library with RepeatMasker 4.1.2 (Smit et al. 1996-2010). The sampled reads were mapped to dimers of satDNA consensus sequences, and for smaller satDNAs, several monomers were concatenated until reaching roughly 150 bp array length. The resulting RepeatMasker .*align* file was then parsed to the script *calcDivergenceFromAlign*.*pl* from RepeatMasker utils. The relative abundance of each satDNA sequence was then estimated as the proportion of nucleotides aligned with the reference sequence with respect to the total Illumina library size.

The RepeatModeler2 library was then merged with the satDNA library produced here and with known avian consensus sequences of transposable elements from Repbase (Bao et al. 2015), Dfam (Storer et al. 2021, 2021), flycatcher, blue-capped cordon-bleu, hooded crow, and paradise crow (Boman et al. 2019; Peona et al. 2021; Suh et al. 2018; Weissensteiner et al. 2020). This library was then used to annotate the genome assembly with RepeatMasker 4.1.2 (Smit et al. 1996-2010). The annotation produced was processed with the script *calcDivergenceFromAlign*.*pl* from RepeatMasker utils to calculate the divergence between repeats and their consensus sequences using the Kimura 2-parameter distance corrected for the presence of CpG sites.

#### Full-length transcript sequencing and genome annotation

We aimed to establish a high-quality genome annotation based on full-length transcripts. To this end, for each of the abovementioned seven tissues, the NGS platform of the University of Berne, Switzerland, prepared an Iso-Seq library using the SMRTbell Express Template Prep Kit 2.0 (Pacific Biosciences). These seven libraries were then sequenced on three separate SMRT cells 8M, sequencing twice two tissues (brain and testis, liver and kidney) and once three tissues (lung, heart, muscle) per SMRT cell. Sequencing of these SMRT cells was conducted on a Pacific Biosciences Sequel II instrument at the Genomic Technologies Facility in Lausanne, Switzerland. As the libraries underloaded, all libraries were jointly sequenced on an additional SMRT cell 8M on a Pacific Biosciences Sequel IIe at the NGS platform of the University of Berne.

Circular consensus sequences (CCS), full-length non-chimeric transcripts, and polished high- and low-quality transcripts were obtained by the NGS platform at the University of Bern using the Isoseq 3 pipeline (ICS v10.1). Polished full-length isoforms for each tissue were then separately mapped to the reference genome using Minimap v2.2 (-ax splice) (Li 2018, 2021). Transcriptome annotations were generated by first collapsing redundant transcripts using TAMA collapse (-x no_cap), before generating open reading frame (ORF) and nonsense-mediated mRNA decay (NMD) predictions using the scripts implemented in TAMA-GO (Kuo et al. 2020) for each of the seven tissues. We then evaluated tissue-specific transcriptome completeness using BUSCO (Simão et al. 2015) with the avian dataset aves_odb10 (8’338 BUSCO) in BUSCO 5.0.0. Additional transcriptome annotation statistics were obtained using the *agat_sp_statistics*.*pl* script implemented in the AGAT toolkit (Dainat 2019).

We annotated the repeat soft-masked genome using GeMoMa version 1.9 (Keilwagen et al. 2018; Keilwagen et al. 2019), a homology-based gene prediction tool. This tool is based on the annotation of protein-coding genes and intron position conservation in a reference genome to predict the annotation of protein-coding genes in the target genome. We used the genomes of chicken (GCA_016699485.1; International Chicken Genome Sequencing Consortium 2004), zebra finch (GCA_003957565.2; Warren et al. 2010), silvereye (GCA_001281735.1; Cornetti et al. 2015), and collared flycatcher (GCA_000247815.2; Ellegren et al. 2012; Kawakami et al. 2014) as references for the homology-based gene prediction, along with the reference individual’s transcriptome obtained from Iso-Seq data to incorporate RNA evidence for the splice prediction. Using the Extract RNA-seq Evidence tool implemented in GeMoMa, we obtained intron position and coverage. This information was fed into the GeMoMa pipeline (GeMoMa.m=200000, AnnotationFinalizer.r=SIMPLE, pc=true, and o=true) to obtain predicted protein-coding gene models. To account for redundancies/duplicates resulting from the predicted protein-coding genes potentially stemming from each of the four reference species, genome annotation completeness was assessed by recomputing BUSCO using the BUSCOrecomputer tool in GeMoMa.

Functional annotation of protein-coding genes was obtained with InterProScan 5.59 (Jones et al. 2014; Paysan-Lafosse et al. 2022). InterProScan ran with the following settings: *-goterms -iprlookup -appl CDD, COILS, Gene3D, HAMAP, MobiDBLite, PANTHER, Pfam, PIRSF, PRINTS, PROSITEPATTERNS, PROSITEPROFILES, SFLD, SMART, SUPERFAMILY, TIGRFAM*).

Predicted protein-coding genes were further annotated through a protein Blast search (-evalue 0.000001, -seg yes, -soft_masking true, -lcase_masking) against the Swiss-Prot database (Uniprot Consortium 2019). We then merged the predicted protein-coding gene models and the functional annotation using the *agat_sp_manage_functional_annotation*.*pl* script, obtained summary statistics using *agat_sp_statistics*.*pl* and *agat_sp_functional_statistics*.*pl*, both implemented in the AGAT toolkit. Gene ontology (GO-terms) were visualised with WEGO 2.0 (wego.genomics.cn).

## Results and Discussion

### Nuclear genome assembly

The polished, unzipped primary assembly contained a total of 1’681 contigs, of which all were >25 kb long and 1’610 were >50 kb long (**Tab. 1**). Total assembly length was 1.29 Gb, with the longest contig spanning 45.3 Mb, contig N50 of 8.6 Mb, and half of the assembly placed in 35 contigs. Avian BUSCO were 96.9 % complete, with 90.6 % being single-copy genes (**Tab. 1**).

**Table 1.** Assembly statistics.

|  |  | Falcon unzip,<br>Arrow | + Pilon,<br>purge_dups | + Tigmint | + 3D DNA<br>(all) | + 3D DNA<br>(chrom) |
| --- | --- | --- | --- | --- | --- | --- |
| Basic stats | No. contigs/scaffolds* | 1'681 | 381 | 383 | 588* | 32* |
|  | No. contigs/scaffolds* > 50 kb | 1'610 | 347 | 348 | 143* | 31* |
|  | Assembly length (Gb) | 1.29 | 1.04 | 1.04 | 1.04* | 1.00* |
|  | Contig/scaffold* N50 (Mb) | 8.6 | 13.5 | 12.6 | 69.6* | 69.7* |
|  | Contig/scaffold* L50 | 35 | 23 | 24 | 6* | 5* |
|  | Largest contig/scaffold* (Mb) | 45.3 | 45.3 | 45.3 | 148.4* | 148.4* |
| BUSCO | Complete (%) | 96.9 | 96.4 | 96.4 | 96.2 | 95.5 |
|  | Complete single-copy (%) | 90.6 | 95.9 | 95.9 | 95.7 | 95.1 |
|  | Complete duplicated (%) | 6.3 | 0.5 | 0.5 | 0.5 | 0.4 |
|  | Fragmented (%) | 0.7 | 0.7 | 0.7 | 0.9 | 0.9 |
|  | Missing (%) | 2.4 | 2.9 | 2.9 | 2.9 | 3.6 |

\* Where numbers concern scaffolds instead of contigs, this is indicated by an asterisk.

Purging duplicated contigs resulted in an assembly constituted of 381 contigs with a total assembly length of 1.04 Gb, contig N50 of 13.5 Mb and half of the assembly placed in 23 contigs (**Tab. 1**). After this step, BUSCO completeness remained at 96.4 %, but an improvement to nearly 96 % single-copy BUSCOs was achieved (**Tab. 1**).

Starting from an already highly contiguous assembly, the linked-read data did not yield any scaffolding. Still, Tigmint detected several supposed mis-assemblies and split the assembly into 451 scaffolds. However, an alignment of the original contigs in D-Genies (Cabanettes and Klopp 2018) showed that all but one of the original contigs (see below) were collinear with the collared flycatcher genome. Given this result and that the proximity ligation data would correct mis-assemblies in subsequent steps, we decided to keep the original contigs except for one aligning to flycatcher chromosomes 2 and 3. For the latter contig, we used the output of Tigmint that split the contig in line with the alignment. The two split parts covered all but 12’527 bp of the original contig. Visual inspection of the missing sequence showed that it almost entirely consisted of repeats. We left this sequence in the assembly as a separate contig.

The proximity ligation information obtained through Hi-C scaffolding corrected a number of scaffolds, resulting in a higher number of scaffolds (588) than the number of contigs it started from (383). However, the scaffolding yielded a highly contiguous chromosome-scale assembly (N50, 69.6 Mb; L50, 6) with BUSCO completeness of still >96 % and almost all BUSCOs in single copy (**Tab. 1**). This final assembly contained all macrochromosomes and the majority of microchromosomes usually found in the latest generation of avian genome assemblies (Kapusta et al. 2017; Peona 2021; Rhie et al. 2021). 96 % of the assembly was placed into chromosome models, and the chromosome-only assembly covered still 95.5 % of BUSCO (**Tab. 1**).

The final assembly length closely matched the one of previous linked-read-based assemblies of the same species and closely related ones (Lutgen et al. 2020; Schweizer et al. 2019). The genome size estimated from the k-mer distribution of linked reads sequence was between 1.105 and 1.106 Gb, with 0.925-0.926 Gb of unique and 0.179-0.180 Gb (16 %) repeat sequence and 0.75-0.76 % heterozygosity (GenomeScope model fit 98-99 %). The full final reference genome assembly thus covered 94 % of the genome size estimate, with 90 % of the estimated genome size placed in chromosomes. 96% of the assembly were placed in 33 chromosomes with homologs in collared flycatcher, zebra finch and chicken, according to which we adapted the chromosome nomenclature. The differences in genome size estimates based on the k-mer approach and the genome assembly length is likely the result of highly repetitive sequences (e.g., centromeres, telomeres, satDNAs) that collapsed during the assembly process (Peona et al. 2018).

### Mitogenome assembly

MitoFinder and MitoVGP assembled mitogenomes of 16’944 bp and 18’631 bp length, respectively. The mitochondrial contigs assembled by the two pipelines were congruent, except for 9 single base pair mismatches, for a 1’827 bp long insert in the MitoVGP assembly and of a 141 bp long insert in the MitoFinder assembly. We decided to not consider either of these inserts in the final mitogenome assembly for the following reasons. First, neither of the inserts was observed in the mitogenomes of Isabelline and Northern Wheatear. For the long insert in the MitoVGP assembly, moreover, the coverage of short-reads mapped to the MitoVGP assembly was strongly reduced (**Fig. S1**), and the insertion constituted a partial duplication of *nd6*, duplications of two tRNAs (Glu, Pro) and a partial duplication of the control region unlikely to be real. The short insert in the MitoFinder assembly was not observed in the other wheatear mitogenomes, and if real, we would expect long reads to cover this insert. Because regarding base calls short reads are expected to have higher quality, we retained the MitoFinder assembly, but without the 141 bp insert as final mitogenome.

The final mitogenome (as also both original assemblies) contained all 13 protein-coding genes, two rRNAs, and 22 tRNAs (**Fig. 2**). All genes, except eight tRNAs and *nd6*, were located on the heavy DNA strand. Both gene order and strandedness were concordant with those observed in Northern Wheatear (*O. oenanthe*) (Wang et al. 2020).

**Figure 2.**
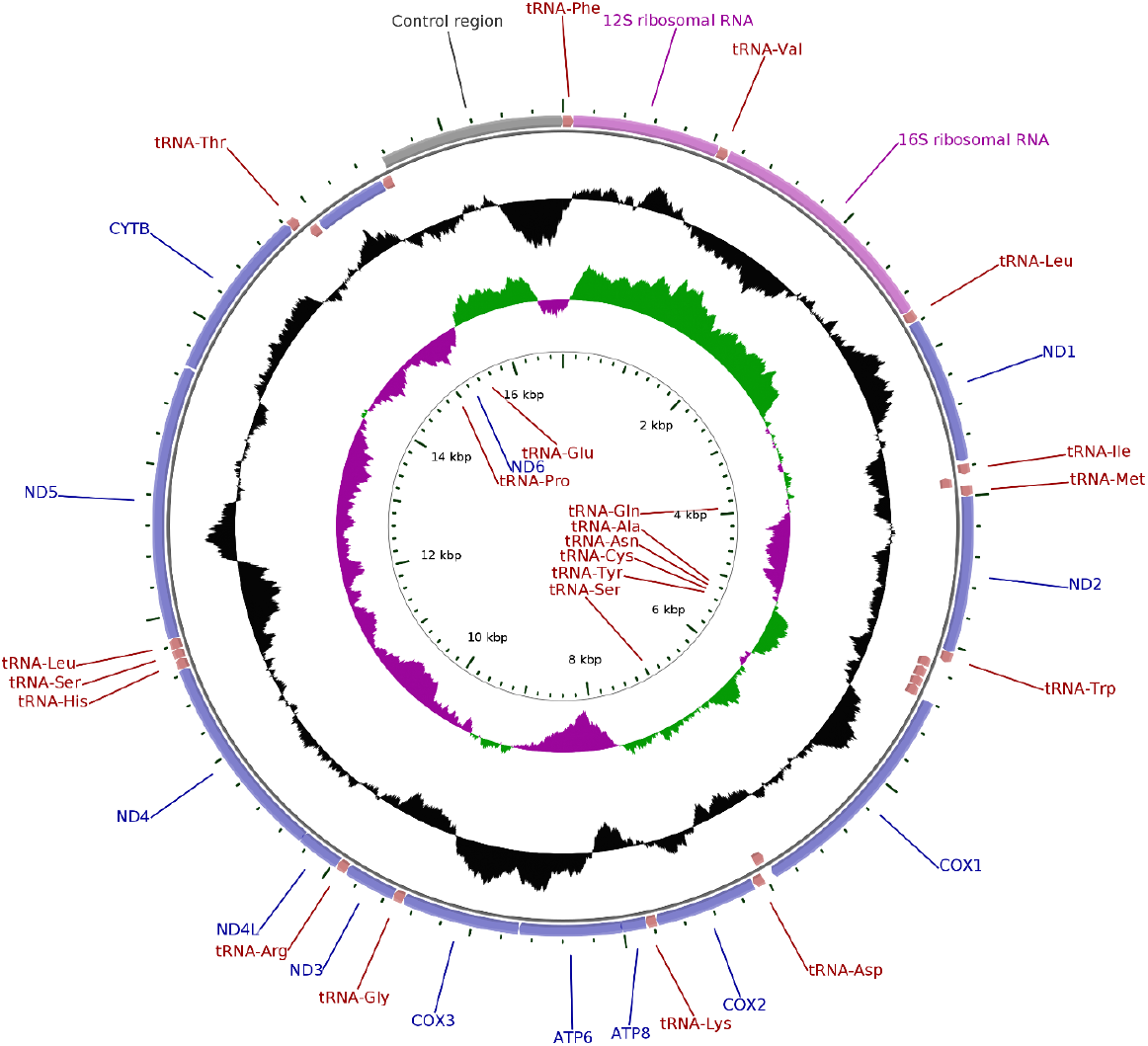
Circular sketch map of the *O. melanoleuca* mitogenome assembly. The outer circle shows coding sequences (purple), rRNAs (pink), and tRNAs (red). The black trace on the middle circle indicates GC content. On the inner circle, positive and negative GC skews in nucleotide composition are indicated by green and magenta, respectively.

### Repetitive element annotation

The *de novo* identification of repetitive elements resulted in the characterisation of 572 raw consensus sequences from RepeatModeler2 and 16 satellite DNA consensus sequences from RepeatExplorer2. The consensus sequences from RepeatModeler2 were filtered from tandem repeats and protein-coding genes. This resulted in a final library of 477 consensus sequences (**File S1**). Among these consensus sequences, RepeatModeler2 classified 226 sequences as LTR retrotransposons, 98 as LINE retrotransposons, 21 as DNA transposons, 5 as SINE retrotransposons, and 112 sequences were unclassified (“unknown”).

The genome assembly annotation run with RepeatMasker using the repeat library produced here and merged with already known avian repeats showed that ∼10% of the assembled genome is repetitive (**Fig. 3A, Tab. S1, File S2**). This finding indicates that many repeats collapsed during the genome assembly process. An example of this were satDNAs that represented ∼0.8% of the sequenced reads but only < 0.3% of the genome assembly, suggesting that satDNA repeats (such as in (peri-)centromeric and (sub-)telomeric regions) are the most collapsed repeats. Most of the repeats annotated were LTR and LINE retrotransposons (**Fig. 3A**). While it is common to find LINEs as most abundant TEs in avian genomes (Galbraith et al. 2021; Kapusta and Suh 2017; Manthey et al. 2018; Peona, Blom et al. 2021), it is less common to find so similar percentages of LINE and LTR retrotransposons. This is especially true for a male genome assembly such as the present one here that does not include the W chromosome which is highly enriched in LTRs and acts as a refugium for most of the full-length genomic LTR elements in birds (Peona et al. 2021; Warmuth et al. 2022). The transposable element landscape (**Fig. 3B**) suggests that LINE retrotransposons experienced a drop in their genomic accumulation in recent times (0-5% divergence; **Fig. 3B**), whereas LTR retrotransposons kept accumulating at the same rate. Such a recent replacement of LINE retrotransposon activity with a diversity of LTR retrotransposons has been noted in other songbirds and seems to have occurred independently in the so far analysed passerine families, i.e., estrildid finches (Warren et al. 2010, Boman et al. 2019), flycatchers (Suh et al. 2018), crows (Weissensteiner et al. 2020), and birds-of-paradise (Peona et al. 2021). Finally, the satDNA landscape (**Fig. 3B**) shows that satDNA arrays experienced differential amplification in copies number in recent times (0-10% divergence), implying fast evolution of this genomic fraction in the genome (Peona et al. 2022).

**Figure 3.**
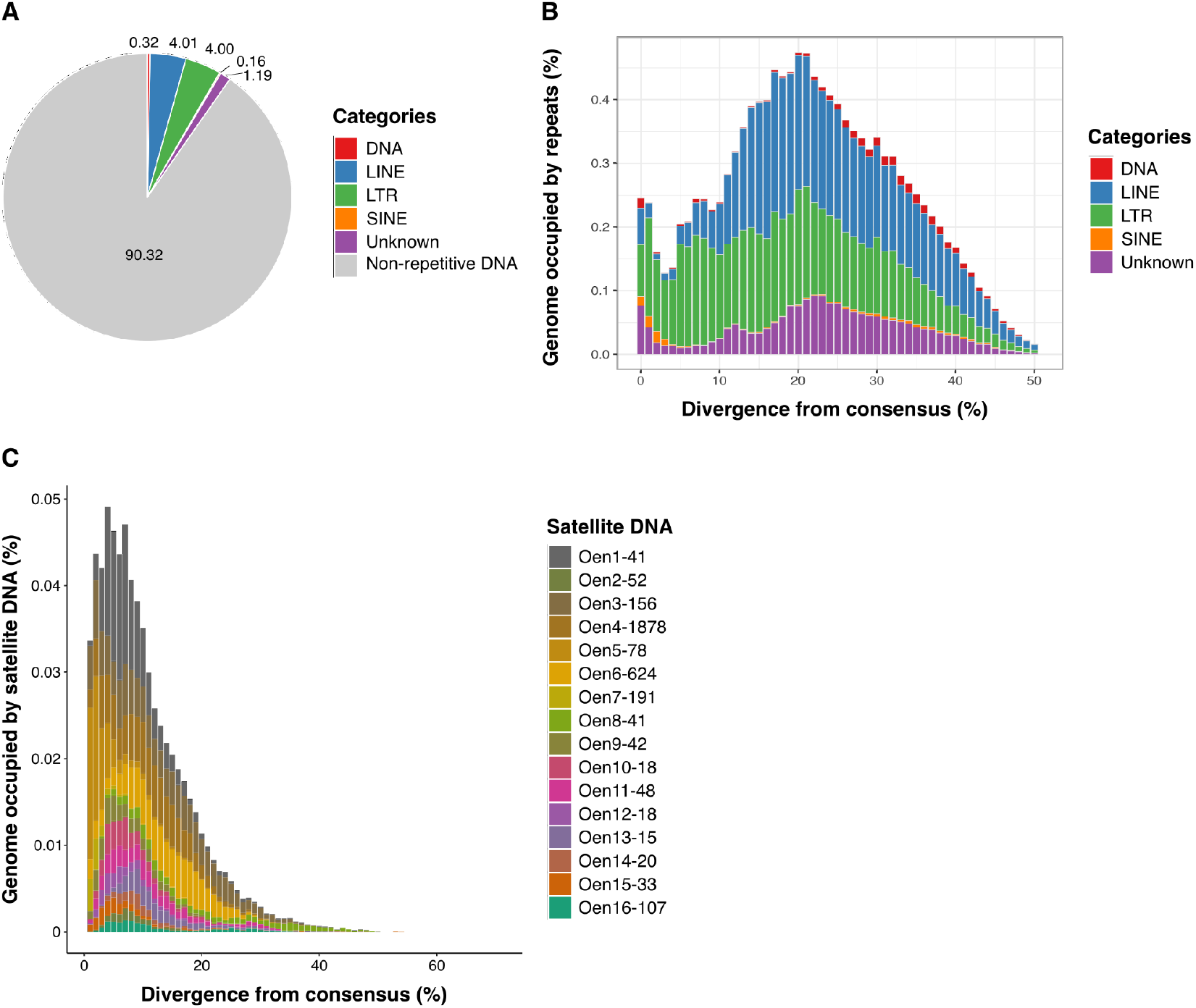
Repeat annotation landscapes. **A**) Pie-chart summarizing the transposable element content annotated in the genome assembly. **B**) Transposable element landscape. The divergence between interspersed repeat copies and their consensus sequences is shown on the X-axis as genetic distance calculated using the Kimura 2-parameter distance. The quantity of the genome assembly occupied by transposable elements in Mb is shown on the Y-axis. **C**) Satellite DNA landscape. The divergence between the satellite DNA consensus sequences and sequences annotated in the short-read library is shown on the X-axis as genetic distance calculated using the Kimura 2-parameter distance. The percentage of the genome (short reads) annotated as satellite DNA is shown on the Y-axis.

### Transcriptome sequencing, genome annotation, and gene function prediction

Iso-Seq sequencing yielded a total of 3’583’686 CCS reads (80’050-1’087’892 reads per tissue, **Tab. 2**). This resulted in numbers of high-quality isoforms ranging from 9’182 to 47’491 per tissue. On average 8’138 genes were predicted per tissue, ranging from 3’375 in muscle to 10’924 in liver. Transcriptome completeness evaluated through BUSCO ranged from 22.80% to 58.10% complete BUSCO per tissue (**Tab. 3**).

**Table 2.** Iso-Seq data characterization and transcriptome completeness.

|  |  | Brain | Heart | Kidney | Liver | Lung | Muscle | Testis |
| --- | --- | --- | --- | --- | --- | --- | --- | --- |
| Transcriptome | No. of CCS reads | 434'902 | 160'998 | 726'466 | 1'087'892 | 658'476 | 80'050 | 284'691 |
|  | High-quality isoforms | 32'188 | 15'937 | 45'097 | 47'491 | 42'209 | 9'182 | 21'311 |
|  | Low-quality isoforms | 353 | 101 | 616 | 384 | 488 | 60 | 154 |
|  | No. of genes | 10'210 | 9'741 | 9'063 | 10'924 | 5'118 | 3'375 | 8'532 |
|  | Mean gene length (bp) | 24910 | 21255 | 16350 | 15125 | 17861 | 16803 | 17120 |
|  | No. of CDS | 26726 | 25442 | 25790 | 27202 | 8946 | 5082 | 18365 |
|  | Mean CDS length (bp) | 1040 | 1062 | 932 | 823 | 904 | 998 | 978 |
|  | No. of exons | 237'491 | 222'791 | 204'320 | 194'325 | 71'818 | 42'504 | 149'404 |
|  | Mean exon length (bp) | 249 | 248 | 223 | 225 | 216 | 221 | 207 |
|  | Mean no. of exons/mRNA | 8.9 | 8.8 | 7.9 | 7.1 | 8.4 | 8.4 | 8.1 |
| BUSCO | Complete (%) | 58.10 | 57.30 | 48.30 | 49.40 | 31.10 | 22.80 | 45.90 |
|  | Single-copy (%) | 40.70 | 40.70 | 33.50 | 34.60 | 27 | 20.50 | 34.20 |
|  | Duplicated (%) | 17.40 | 16.60 | 14.80 | 14.80 | 4.10 | 2.30 | 11.70 |
|  | Fragmented (%) | 2.20 | 2.10 | 2.60 | 3.20 | 1.60 | 0.80 | 2.00 |
|  | Missing (%) | 39.70 | 40.60 | 49.10 | 47.40 | 67.30 | 76.40 | 52.10 |

**Table 3.**
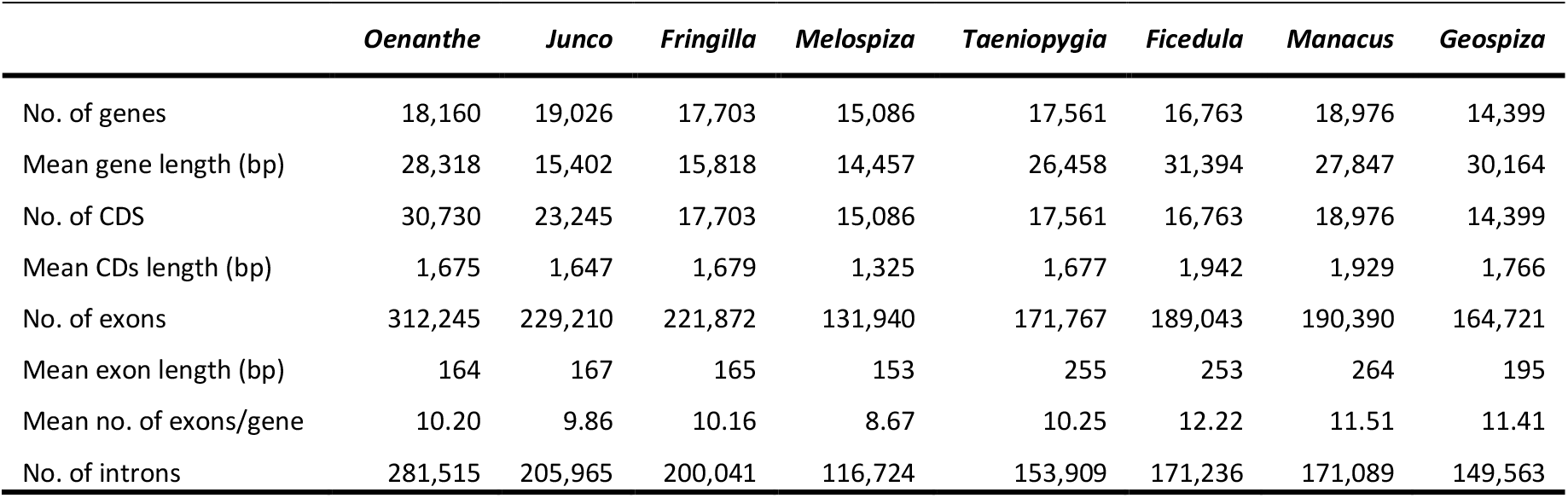
Summary statistics for the predicted gene annotation and comparison to other bird species (*Oenanthe melanoleuca* versus *Junco hyemalis, Fringilla coelebs, Melospiza melodia, Taeniopygia guttata, Ficedula albicollis, Manacus vitellinus*, and *Geospiza fortis*). Modified from Friis et al. (2022).

|  | *Oenanthe* | *Junco* | *Fringilla* | *Melospiza* | *Taeniopygia* | *Ficedula* | *Manacus* | *Geospiza* |
| --- | --- | --- | --- | --- | --- | --- | --- | --- |
| No. of genes | 18,160 | 19,026 | 17,703 | 15,086 | 17,561 | 16,763 | 18,976 | 14,399 |
| Mean gene length (bp) | 28,318 | 15,402 | 15,818 | 14,457 | 26,458 | 31,394 | 27,847 | 30,164 |
| No. of CDS | 30,730 | 23,245 | 17,703 | 15,086 | 17,561 | 16,763 | 18,976 | 14,399 |
| Mean CDs length (bp) | 1,675 | 1,647 | 1,679 | 1,325 | 1,677 | 1,942 | 1,929 | 1,766 |
| No. of exons | 312,245 | 229,210 | 221,872 | 131,940 | 171,767 | 189,043 | 190,390 | 164,721 |
| Mean exon length (bp) | 164 | 167 | 165 | 153 | 255 | 253 | 264 | 195 |
| Mean no. of exons/gene | 10.20 | 9.86 | 10.16 | 8.67 | 10.25 | 12.22 | 11.51 | 11.41 |
| No. of introns | 281,515 | 205,965 | 200,041 | 116,724 | 153,909 | 171,236 | 171,089 | 149,563 |

The Iso-Seq transcriptomes were then used as splice evidence in GeMoMa to perform a predominantly homology-based annotation of the reference genome. We predicted 18’160 protein-coding genes with a total of 312’245 exons and 281’515 introns. The number of exons, CDS, and introns was higher for our *O. melanoleuca* annotation compared to the annotations of other songbirds, such as *Junco cyemalis, Fringilla coelebs, Melospiza melodia, Taeniopygia guttata, Manacus vitellinus and Geospiza fortis* (**Tab. 3**). Mean gene length, CDS length, exon length, and number of exons per gene, on the other hand, were in the range of values obtained for the abovementioned songbird annotations (**Tab. 3**). 17’566 (96%) of the 18’160 predicted genes were annotated with protein families or function assignment. 12’471 (69%) genes obtained a GO term assignment through InterProScan. The most abundant GO terms were associated with “cell part”, “cell” and “membrane” in the cellular component category, “binding” in the molecular function category and “cellular metabolic process” or “metabolic process” in the biological process category (**Fig. S2**). BUSCO completeness of the final annotation as judged from avian BUSCO (n=8’338) was 98%, with 97.4% single copy BUSCO, 0.6% duplicated BUSCO, 0.6% fragmented BUSCO, and 1.5% missing BUSCO. This suggests an accurate and rather complete annotation.

## Data Availability

All data, including the assembly, its annotation, and the original sequencing data are available on the European Nucleotide Archive under project assession *XY (to be provided upon acceptance)*. Code for the repeat analysis is available on https://github.com/ValentinaBoP/WheatearGenome Analysis.

Supplementary material is available on figshare.com.

## Acknowledgements

We warmly thank Marta Burri for preparing RNA, and Giulio Formenti, Remi Allio, and Lauren Coombe for support with computational questions. We are indebted to NGI Uppsala, namely Mai-Britt Mosbech and Olga Vinnere Pettersson for UHMW DNA extraction and long-read sequencing and Ignas Bunikis for running the primary assembly, as well as NGI Stockholm for the preparation of linked-read and Hi-C sequencing data. Finally, we thank the NGS platform at the University of Berne, namely Pamela Nicholson and Catia Coito for the preparation of Iso-Seq data. Computations were performed on resources provided by the Swedish National Infrastructure for Computing (SNIC) through the Uppsala Multidisciplinary Center for Advanced Computational Science (UPPMAX) and the High-Performance Computing Cluster EVE, a joint effort of the Helmholtz Centre for Environmental Research (UFZ) and the German Centre for Integrative Biodiversity Research (iDiv) Halle-Jena-Leipzig. We thank the administration and support staff of EVE: Thomas Schnicke and Ben Langenberg (UFZ), and Christian Krause (iDiv).

## Conflict of Interest

The authors declare no conflict of interest.

## Funder Information

The present project was supported by German Research Foundation (DFG), grant number BU3456/3-1 to RB and the National Research Fund (FNR) Luxembourg, grant number 14575729, to DL; OMPG was supported by the Swedish Research Council Vetenskapsrådet (grant number 2020-03866); VP was supported via grants to AS from the Swedish Research Council Vetenskapsrådet (grant number 2020-04436) and the Swedish Research Council Formas (2017-01597). NAK was supported by a Georg Foster Research Stipend of the Alexander von Humboldt Foundation.

